# Parallel visual processing in the suprachiasmatic region of a diurnal mammal

**DOI:** 10.64898/2026.09.07.749833

**Authors:** Patrycja Orlowska-Feuer, Jessica Rodgers, Rose Richardson, Asshen Dedigama Acharige, Martha Davey, Nina Milosavljevic, Annette E. Allen, Timothy M. Brown, Frank P. Martial, Riccardo Storchi, Beatriz Bano-Otalora, Robert J. Lucas

## Abstract

Light shapes mammalian physiology and behaviour through hypothalamic pathways, yet how these pathways process visual information in day-active species remains poorly understood. Based largely on work in nocturnal rodents, the suprachiasmatic nucleus (SCN) is classically viewed as a retinal target that encodes ambient illumination for circadian entrainment. We combine anatomical tracing, light-evoked c-Fos mapping and high-density silicon-probe electrophysiology to define visual processing in the suprachiasmatic region of the diurnal murid *Rhabdomys pumilio*. We find that retinal input and light-evoked activity extend beyond the SCN into rostral and lateral peri-SCN hypothalamus. Within the SCN, neurons encode irradiance with graded changes in maintained firing broadly similar to those observed in parallel recordings of nocturnal *Mus domesticus*. Across the wider suprachiasmatic region, *Rhabdomys* neurons show selectivity for more dynamic visual stimuli, including luminance transitions, contrast and temporal structure. Unsupervised clustering identifies five functionally distinct hypothalamic visual response classes with different anatomical distributions. These findings reveal an expanded hypothalamic visual network in a diurnal mammal, placing SCN irradiance coding within a broader system for processing dynamic visual features.

## Introduction

Light exerts widespread effects on mammalian physiology and behaviour, from setting circadian phase (entrainment) and timing sleep to driving changes in alertness, mood, metabolism, and neuroendocrine state (1, 2). A principal anatomical route for these effects is the retinohypothalamic tract (RHT), which conveys retinal signals to the anterior hypothalamus, particularly the suprachiasmatic nucleus (SCN), site of the principal circadian pacemaker and origin of daily variations in physiology and behaviour (3). In nocturnal rodents, extensive electrophysiological work has shown that SCN neurons respond robustly to light (4–8) to encode stimulus irradiance across a wide dynamic range, providing a neural representation of ambient illumination (7–13).

The daily light experience of day-active animals differs markedly from nocturnal species, which are primarily exposed to light around twilight (14–16). Yet neurophysiological studies of SCN light responses in diurnal species remain sparse. Recordings from squirrels and *degus* confirm that electrophysiological responses to light can be readily recorded in the day-active SCN (17, 18), however two important questions remain unresolved. First, is the SCN visual response different in nocturnal vs diurnal species? The relatively small number of neurons from diurnal species whose light responses have been recorded more often decreased firing than is typically the case in nocturnal rodents (7, 8, 13), leading to the interesting hypothesis that light may be primarily inhibitory for the diurnal SCN (17, 18). Second, is the expansion of visual information capacity that is a hallmark of diurnality (19, 20) apparent in the visual hypothalamus? SCN size does not show large inter-species variability, but a recent fMRI study in humans raises the possibility that the neighbouring anterior hypothalamus may be more visually responsive than expected based upon recordings in nocturnal rodents (21).

Here we set out to address these knowledge gaps in suprachiasmatic visual processing in diurnal species by employing the four striped African mouse (*Rhabdomys pumilio)*, a diurnal murid rodent (22–25). *Rhabdomys* offers a close phylogenetic comparison with the nocturnal mouse (*Mus domesticus)* and rat *(Rattus norvegicus)* while having an eye, retina and visual brain with canonical features of diurnal species (26–28). The RHT is intact in *Rhabdomys* (29) and a recent *ex vivo* study showed primarily inhibition following electrical stimulation targeting the optic nerve (30). However, there has been no characterisation of light-evoked electrophysiological activity in the suprachiasmatic hypothalamus in this species. Here, we describe retinal input at both anatomical and functional levels in *Rhabdomys*. We find that visual responsiveness extends beyond the SCN to an extent quite distinct from related nocturnal species. This anatomical distribution is mirrored by functional diversity, with hypothalamic neurons segregating into five functionally distinct visual pathways encoding a diversity of visual features beyond the expected ability to report ambient light. Our data thus reveal a diverse and distributed suprachiasmatic visual centre in the diurnal hypothalamus.

## Results

### Widespread visual responses across the suprachiasmatic region of *Rhabdomys pumilio*

We started by revisiting the retinal input to the anterior hypothalamus of *Rhabdomys* (29) by combining anterograde tracer cholera toxin b-subunit (CTb) with iDISCO brain clearing and light-sheet microscopy to obtain 3D representation of this projection. We found CTb staining primarily contralateral to the injected eye within an ellipse bounded by the ventral surface and 3rd ventricle above the chiasm (matching the location of the SCN (31)), but sparse fibres extended to more lateral regions rostral and parallel to the main locus of projection (Fig. 1A; Movie 1 and 2).

**Fig. 1.**
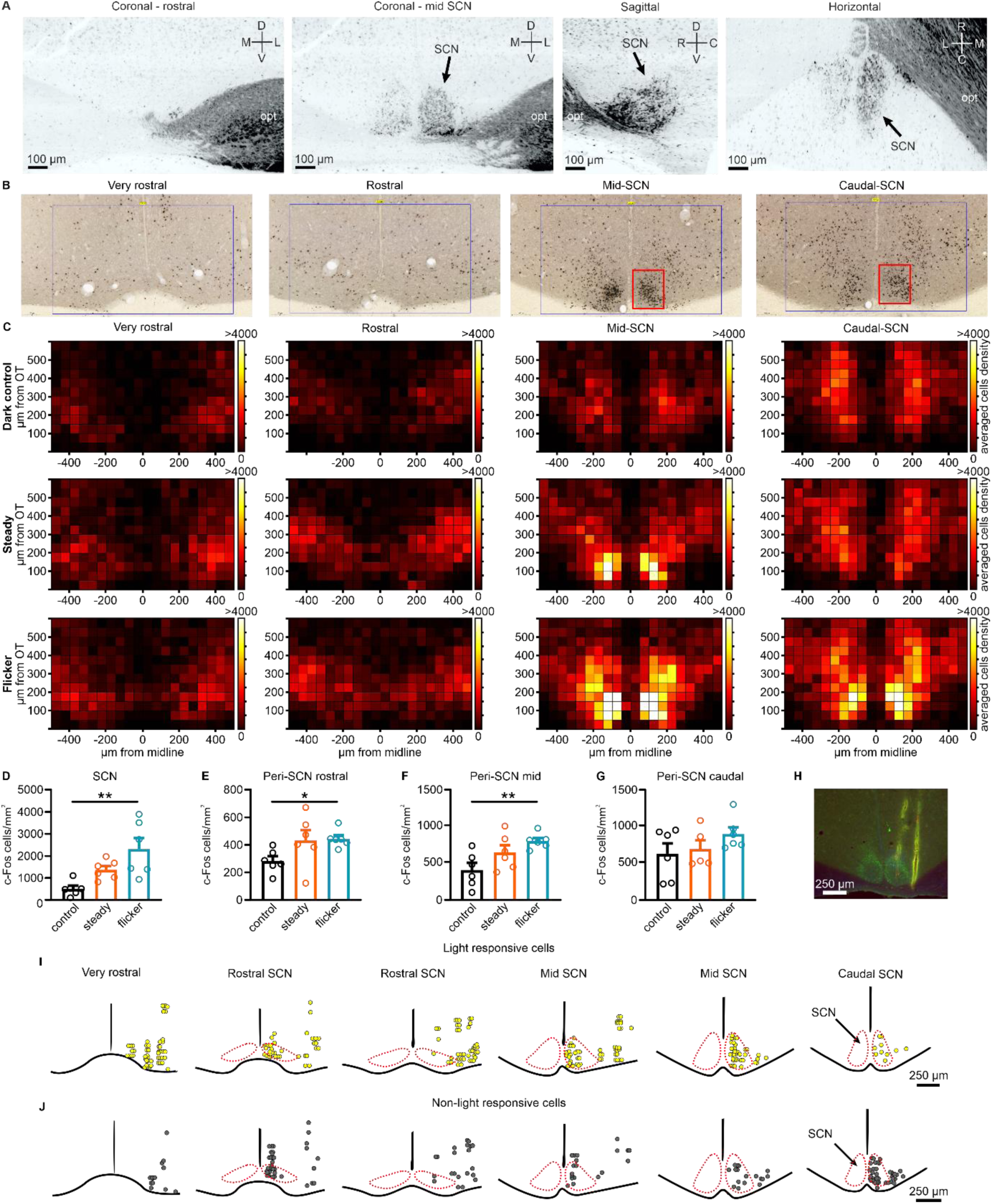
Retinal input and light-evoked activation extend beyond the SCN in diurnal *Rhabdomys.* A) CTb-labelled retinal fibres in the anterior hypothalamus following unilateral intravitreal injection in *Rhabdomys pumilio*. Coronal sections are shown rostral to the SCN and at the mid-SCN level, with sagittal and horizontal views through the mid-SCN. Images were generated from 3D volume reconstructions acquired by light-sheet microscopy after iDISCO brain clearing (Movie 2). Crosses indicate image orientation. B) Representative coronal sections showing c-Fos immunoreactivity in the anterior hypothalamus after 30 min exposure to 4 Hz flicker light at ZT16. Dark nuclear staining indicates c-Fos-positive cells. Sections correspond to the rostro-caudal levels mapped in C. Blue boxes indicate the full anterior hypothalamic regions used for c-Fos mapping (1000 µm × 600 µm), whereas red boxes indicate anatomically defined SCN regions used for SCN-specific counts. Peri-SCN counts were derived from regions outside the red SCN boxes within the corresponding rostral or lateral anterior hypothalamic sampling areas. C) Heat maps showing the density of c-Fos-positive cells across the anterior hypothalamus in *Rhabdomys*, arranged from rostral to caudal sections. Rows indicate animals kept in darkness, exposed to steady light or exposed to 4 Hz flicker light. D-G) Quantification of c-Fos-positive cells in the SCN and peri-SCN regions across rostro-caudal levels. D) Light exposure produced strong activation in the SCN (one-way ANOVA, p = 0.0042, F(2,15) = 8.083; Tukey multiple-comparison test, control vs flicker: p = 0.0030). E) Flicker light also produced weaker but significant activation in rostral peri-SCN regions (Kruskal–Wallis test, p = 0.0231, KWS = 6.982; Dunn’s multiple-comparison test, control vs flicker: p = 0.0449). F, G) Lateral peri-SCN activation was significant at mid-SCN levels F) but not caudal-SCN levels G) (mid peri-SCN: one-way ANOVA, p = 0.0134, F(2,15) = 5.833; Tukey multiple-comparison test, control vs flicker: p = 0.0107; caudal peri-SCN: one-way ANOVA, p = 0.2623, F(2,14) = 1.475). H) Histological verification of electrophysiological recording sites in *Rhabdomys*. CM-DiI-labelled probe tracks are shown in yellow, and AVP immunostaining is shown in green to define the SCN boundary. I, J) Sequential coronal schematics through the anterior hypothalamus, arranged from rostral to caudal, showing the estimated locations of recorded I) light-responsive units and J) non-light-responsive units in *Rhabdomys*. Coronal drawings were based on the stereotaxic mouse atlas and aligned to species-specific histological landmarks. *p < 0.05, **p < 0.01. L, lateral; M, medial; R, rostral; C, caudal; D, dorsal; V, ventral; opt, optic tract; SCN, suprachiasmatic nucleus.

To aid interpretation of this pattern, we next mapped excitatory responses to light in the suprachiasmatic region using c-Fos as a marker. We processed sections encompassing the SCN and surrounding areas for c-Fos immunocytochemistry from animals held in darkness (control) or presented with 30 min of either steady or 4 Hz flicker light (32) at ZT16 (representative sections are shown in Fig. 1B). Visual inspection of anatomical projections of mean c-Fos positive cell density across the anterior hypothalamus region in each group suggested strong light induction in the region corresponding to the expected location of the SCN (Fig. 1C) and weaker increases across more rostral and, potentially, more lateral regions (termed peri-SCN). We undertook statistical analysis of c-Fos positive neurons in the SCN and in rostral and lateral peri-SCN regions separately, given their difference in anatomical location and relationship with retinal input to this part of the brain. Comparison of c-Fos expression between steady, flicker and control groups showed a significant light effect in SCN especially for the flicker (Fig. 1D). Beyond the SCN, flicker light produced a significant increase in c-Fos expression in sections rostral to the SCN (Fig. 1E). The lateral peri-SCN region showed significant c-Fos induction at the level of mid-(Fig. 1F) but not caudal-SCN (Fig. 1G).

The CTb tracing and c-Fos excitation map confirm retinal projections are associated with a net excitatory response to light in the *Rhabdomys* SCN and neighbouring regions of the anterior hypothalamus. To provide a more comprehensive description of light responses in this region, we turned to *in vivo* electrophysiology. We targeted the anterior hypothalamus with multi-channel extracellular recording electrodes under urethane anaesthesia and started with recording responses to 10 s full-field bright light pulses (5789 melanopic lux EDI). For comparison we extended a published SCN dataset on laboratory mice (*Mus domesticus*) using the same stimulus (13) with further hypothalamic recordings.

Anatomical assignment of recorded units based on stereotaxic coordinates, probe trajectory, probe depth, shank geometry and CM-DiI-labelled electrode tracks (Fig. 1H) revealed recording sites covering the suprachiasmatic region in both species (Fig. 1I&J, Supplementary Fig. 1A&B). We related these to the location of the SCN by counterstaining with AVP and/or DAPI to delineate the boundaries of this nucleus (Fig. 1H). This revealed a striking species difference. In *Rhabdomys*, light-responsive units were not restricted to the SCN but were abundant rostral, dorsal and lateral to the SCN (Fig. 1I&J). By comparison, light responses were rare outside of the SCN in *Mus* (Supplementary Fig. 1A&B).

### Responses of *Rhabdomys* SCN to sustained illumination

For more detailed analysis of light response properties, we focused first on units within the boundaries of the SCN. The population mean response to light across 83 (of 147 recorded over 22 placements from 11 animals) light-responsive units in *Rhabdomys* SCN was qualitatively similar to that in *Mus* (101 of 167 recorded over 19 placements from 9 animals), comprising a transient spike in firing at light on followed by sustained excitation lasting throughout the pulse (Fig. 2A). This, though, masked significant variation at the single-unit level, with examples of units with light-induced reductions in firing as well as more transient responses to the appearance and, in some cases, disappearance of light (Fig. 2B-D). We captured this variability in response polarity as polar plots in which each light-responsive unit was described by a vector whose direction was defined by the change in firing at light ON and OFF and whose magnitude reflected response amplitude (Fig. 2E). In both *Rhabdomys* and *Mus*, there was a clear bias in favour of excitatory responses to light (Fig. 2E), with no significant difference in response polarity between species (Kuiper test, p = 0.100). Calculation of a ‘sustainedness index’, as described for mice (33), revealed that *Rhabdomys* responses were more biased towards the acute response component (two-sample Kolmogorov–Smirnov test, p = 0.0026, D = 0.2646), although both species had units whose response was sustained across the light step (Fig. 2F). Across all types, response latency was lower in *Rhabdomys* (Fig. 2G), in line with previous retinal and visual thalamus recordings of light-responsive cells in this species (27, 28).

**Fig. 2.**
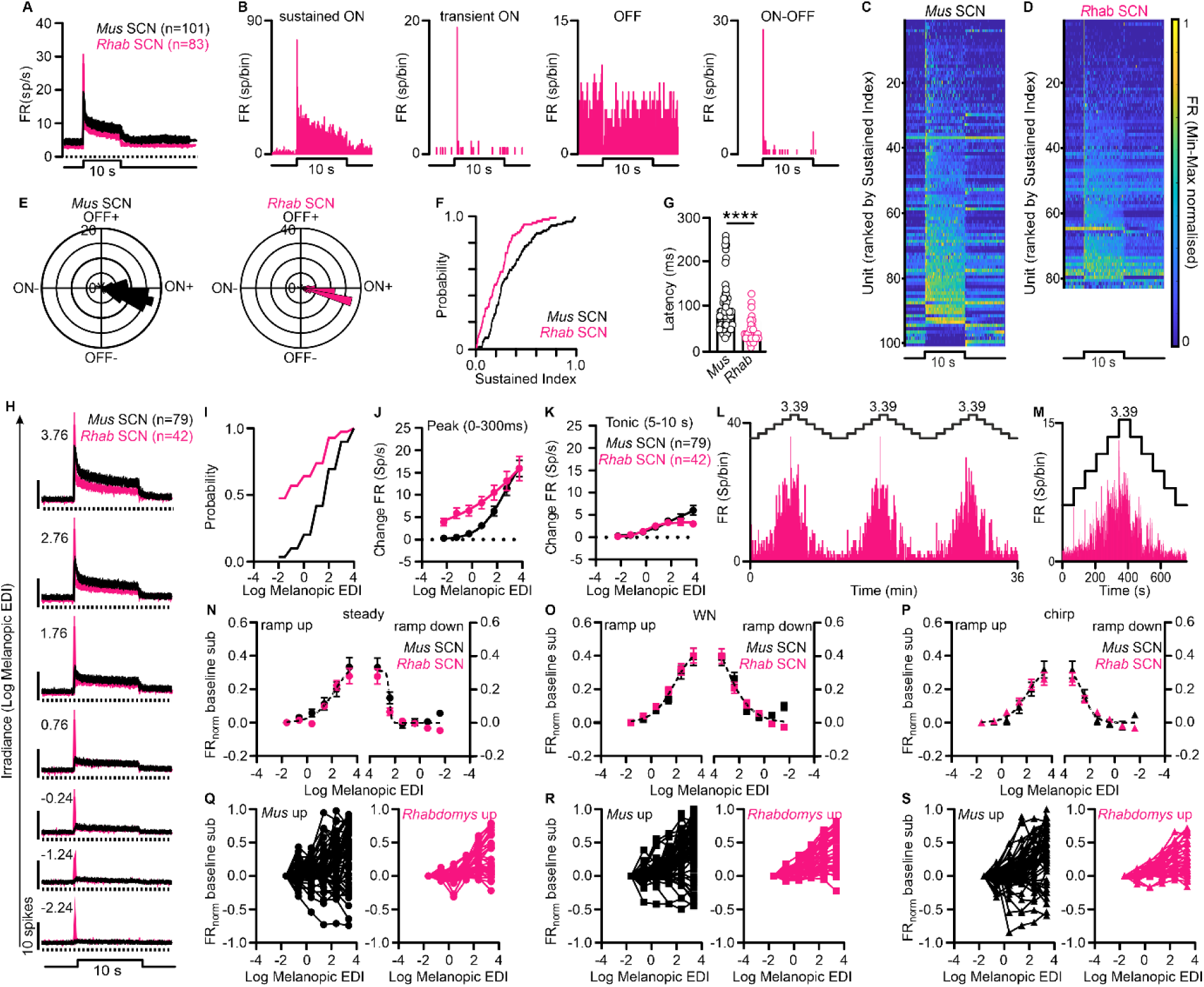
Diurnal *Rhabdomys* SCN neurons show conserved irradiance coding with faster and more transient light responses than *Mus*. A) Mean PSTHs ± SEM (bin size = 0.1 s) in response to a 10 s white-light pulse (5789 melanopic lux EDI) for all light-responsive SCN units in *Mus* (black, n = 101) and *Rhabdomys* (pink, n = 83). B) Example PSTHs (bin size = 0.1 ms) from four representative light-responsive SCN units in *Rhabdomys*. C, D) Heat maps of responses to the 10 s light pulse for all light-responsive SCN units in C) *Mus* and D) *Rhabdomys*, ranked by sustainedness index, with the most sustained responses at the bottom. E) Polar plots summarising response polarity and amplitude show no significant species difference (Kuiper test, p = 0.100). F) Cumulative distribution functions comparing sustainedness index distributions between species (two-sample Kolmogorov–Smirnov test, p = 0.0026). G) Light-response latency for units excited by light onset differs between species (Mann-Whitney test, p < 0.0001, U = 1047). H) Mean PSTHs ± SEM (bin size = 0.1 s) in response to 10 s light pulses of increasing irradiance, arranged from bottom to top, for light-excited SCN cells in *Rhabdomys* (pink, n = 42) and *Mus* (black, n = 79). I) Cumulative distribution of light sensitivity based on responses shown in H. J, K) Irradiance-response relationships for peak firing-rate changes J) during the first 300 ms after light onset (F test, p < 0.0001, F = 11.27) and K) between 5 and 10 s after light onset (F test, p = 0.0225, F = 3.209). L) Spike-rate histogram for an example SCN unit responding to the staircase stimulus, with the stimulus profile shown above. M) PSTH for the same example unit (bin size = 3 s). N–P) Mean baseline-subtracted, normalised firing rate ± SEM of light-responsive cells during ramp-up and ramp-down epochs as a function of irradiance during N) steady light, O) temporal white noise and P) chirp presentation. F-test results are shown for ramp-up and ramp-down fits in each condition (steady light: ramp up: p = 0.3392, F = 1.123, ramp down: p = 0.1524, F = 1.767; WN: ramp up: p = 0.5774, F = 0.6592, ramp down: p = 0.9032, F = 0.1901; chirp: ramp up: p = 0.3603, F = 1.072, ramp down: p = 0.3395, F = 1.122). Q–S) Individual-cell ramp-up irradiance-response traces corresponding to N–P for *Mus* and *Rhabdomys*.

In *Mus*, individual light-responsive SCN cells can respond to ipsi-and/or contralateral retinal stimulation consistent with the bilateral nature of retinal projections in that species (10). We addressed this in *Rhabdomys* in a subset of recordings and found a preponderance of responses to contralateral only stimulation (n = 14/21), consistent with the contralateral bias of retinal input in this species (Fig. 1A and (29)). Bilateral responses were also encountered (n = 7/21) but no unit responded only to ipsilateral stimulation (Supplementary Fig. 2).

The accepted function of retinal input to the mammalian SCN is to encode scene irradiance as a proxy for time of day. We next probed such irradiance coding in a subset of recordings (12 placements in 6 animals) by presenting extended feature-less light pulses (10 s) over a range of intensities (0.005 to 5789 melanopic lux EDI, Fig. 2H). *Rhabdomys* SCN showed high sensitivity to this stimulus, with the acute component of light responses (0-300 ms after stimulus presentation) apparent at the dimmest light tested in almost 50% of excited light-responsive units (n = 20/42 of all excited light-responsive units, Fig. 2I), and the intensity response relationship of this response component showing a significant difference from that of *Mus* (Fig. 2J). As any irradiance coding would be captured by the more sustained response, we next analysed firing 5-10 s after stimulus onset. This tonic component showed irradiance dependence in both species (Fig. 2K). A direct comparison revealed a species difference in irradiance dependence of the tonic component with *Rhabdomys* showing smaller responses at high irradiances than *Mus* (Fig. 2K).

As the step response properties were consistent with irradiance coding in *Rhabdomys* SCN, we continued to probe this behaviour with a more naturalistic ‘staircase’ stimulus (Fig. 2L; (13), comprising 3-8 repeats of ascending+descending epochs of full-field light (0.024 to 2449 melanopic lux EDI), comprised of 30 s steady light followed by 36 s with superimposed higher frequency modulations (temporal white noise (WN) and frequency/contrast chirps). Firing rate for an exemplar *Rhabdomys* SCN neuron across presentations of the staircase irradiance stimulus is shown in Fig. 2L followed by a PSTH for that neuron in Fig. 2M. At the population level (pooled data for all light-responsive neurons) SCN tracked stimulus irradiance with a monotonic increase of firing under all three conditions tested (steady light, WN and chirp), and no differences were observed between species (Fig. 2N-P). Accordingly, at the single cell level light-responsive SCN units in both species mostly showed higher firing at higher irradiances, but there were some units whose firing decreased (13/57 and 6/42 in *Mus* and *Rhabdomys*, respectively; Fig. 2Q-S).

### *Rhabdomys* peri-SCN shows limited sensitivity to ambient light

The most unexpected feature of the *Rhabdomys* dataset was the extent of photosensitivity outside the SCN (significant light responses in 135/215 recorded units, Fig. 1I-J). This has not been reported in studies of nocturnal rodents to our knowledge and was not a feature of our own parallel recordings in mice where photosensitivity in these regions was rare (30/216 units) and mostly at borders of the SCN (Supplementary Fig. 1A&B). Analysis of light-responsive units in *Rhabdomys* peri-SCN revealed that they differ from those within the SCN boundaries in both spontaneous and light-evoked activity (Fig. 3A-B). Light-responsive units within the SCN in *Rhabdomys* showed a progressive reduction in dark activity across the early subjective night (ZT13-17; Fig. 3A), consistent with the intrinsic circadian rhythm in spontaneous activity in this nucleus (34). No such variation in spontaneous activity in the dark was observed in light-responsive peri-SCN units (Fig. 3A), indicating that they are not components of the circadian oscillator (F test comparing linear regression slopes, p < 0.0001). Similarly, while the population mean response to the 10 s pulse within the SCN was a sustained increase in firing, in the peri-SCN it was dominated by transient peaks in firing at light onset and offset (Fig. 3B). There was no consistent increase in response latency in the peri-SCN as might be expected if these were second order to visual responses in the SCN or any other visual centre (Mann-Whitney test, p = 0.1369, Fig. 3C).

**Fig. 3.**
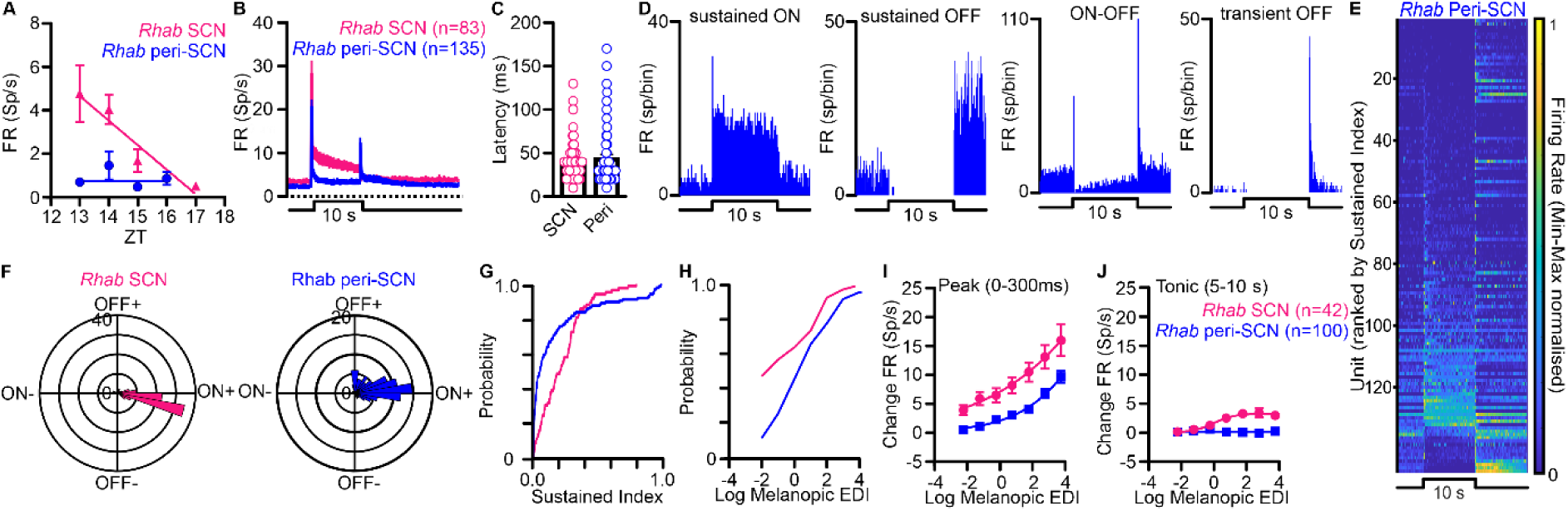
Peri-SCN neurons in diurnal *Rhabdomys* show distinct light-response properties and limited irradiance coding. A) Mean basal firing rate recorded in darkness across ZT13–17 for light-responsive cells in the SCN and peri-SCN. Dark firing rate varied with circadian time in the SCN but not in the peri-SCN, with a significant difference between linear regression slopes (F test, p < 0.0001, F = 28.68). B) Mean PSTHs ± SEM (bin size = 0.1 s) in response to a 10 s white-light pulse (5789 melanopic lux EDI) for all light-responsive SCN cells (pink, n = 83) and peri-SCN cells (blue, n = 135) in *Rhabdomys*. C) Light-response latency for cells excited by light onset did not differ significantly between SCN and peri-SCN populations (Mann-Whitney test, p = 0.1369, U = 4302). D) Example PSTHs (bin size = 0.1 ms) from four representative light-responsive peri-SCN cells in *Rhabdomys*. E) Heat map of responses to the 10 s light pulse for all light-responsive peri-SCN cells, ranked by sustainedness index, with the most sustained responses at the bottom. F) Polar plots summarising response polarity and amplitude reveal a significant difference between SCN and peri-SCN cells (Kuiper test, p = 0.001, K = 5309). G) Cumulative distribution functions comparing sustainedness index distributions between SCN and peri-SCN cells (two-sample Kolmogorov–Smirnov test, p < 0.0001, D = 0.3516). H) Cumulative distribution of light sensitivity for light-responsive SCN and peri-SCN cells in *Rhabdomys*. I, J) Irradiance-response relationships for peak firing-rate changes I) during the first 300 ms after light onset (F test, p < 0.0001, F = 32.80) and J) between 5 and 10 s after light onset (F test, p < 0.0001, F = 41.71).

At the single-unit level, peri-SCN responses to the light step showed substantial diversity in response profile and polarity (shown for exemplar units in Fig. 3D and the population in Fig. 3E). Comparison of polar plots revealed a significant difference in response polarity compared with the SCN (Kuiper test, p = 0.001), which appeared to reflect a greater preponderance of units excited by light offset (Fig. 3F). Peri-SCN light responses were also less sustained and more biased towards responses to light transitions (two-sample Kolmogorov–Smirnov test, p < 0.0001, D = 0.3516; Fig. 3G). When challenged with light steps at increasing irradiance, light-responsive peri-SCN units lacked the very high sensitivity of the *Rhabdomys* SCN transient peak at light onset (Fig. 3H&I; F test, p < 0.0001, F = 32.80) and had little tonic firing at any irradiance (Fig. 3J; F test, p < 0.0001, F = 41.71). By contrast, the few light-responsive cells in *Mus* peri-SCN had response characteristics more like those in the SCN (Supplementary Fig. 3).

### Functional diversity of visual responses

Light pulse responses thus indicate that neurons in the *Rhabdomys* suprachiasmatic region (SCN and peri-SCN) respond to both steady light intensity and abrupt changes in light and that, on average, the weighting of each differs between neurons within *vs* beyond the SCN. We finally set out to gain a more complete understanding of the visual response properties of this neuronal population by presenting a more dynamic stimulus. We employed a full field chirp stimulus comprised of a 3 s light step from dark followed by sinusoidal modulations of increasing temporal frequency from 1 to 8 Hz at 1 Hz/s speed at 97% Michelson contrast and sinusoidal contrast modulation at 2 Hz increasing from 3% to 97% contrast. This chirp has previously been used for objective clustering of visual response types in both *Mus* (retina and visual thalamus) and *Rhabdomys* (visual thalamus) (28, 33, 35) allowing the possibility not only of describing response properties at single unit level but also for functional clusters. We presented this stimulus at a background irradiance of 244.9 melanopic lux EDI to a subset of *Rhabdomys* and *Mus* (n = 7 and 9 respectively) and applied unsupervised clustering to all units showing reproducible responses (based on trial-to-trial correlation). We found that only 12% of units in suprachiasmatic region of *Mus* were classified as responsive to this stimulus (SCN: 33/167 and peri-SCN: 9/187), while 36% (SCN: 61/144 and peri-SCN: 23/80) met this criterion in *Rhabdomys*.

Unsupervised clustering returned 5 functional groups based on chirp response properties (Fig. 4A; methods). These functional clusters appeared to differ in anatomical distribution (Fig. 4B). In *Rhabdomys*, 3 clusters were predominantly within or very close to the SCN (clusters 1, 2 and 4). Conversely, cluster 5 units were primarily located in a region rostral to the SCN. Only cluster 3 had good representation both within and outside (lateral to) the SCN. This distribution was partially recapitulated in mice (Supplementary Fig. 4, Supplementary Table 1). Thus, clusters 3 and 4 were also represented in the *Mus* SCN. On the other hand, clusters 1 and 5 were absent in Mus and the few extra-SCN units in this species fell into cluster 2.

**Fig. 4.**
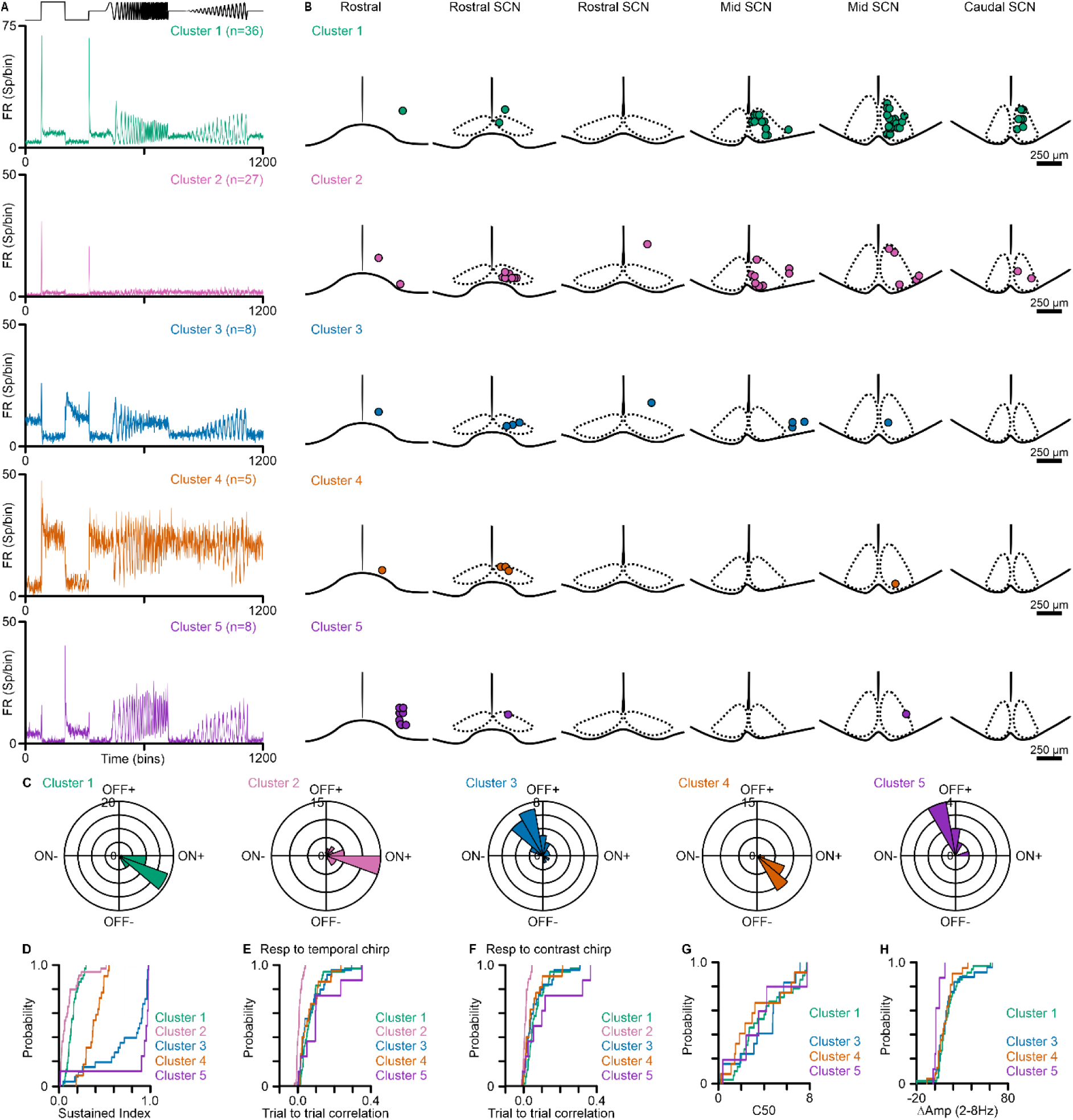
Functional clustering identifies distinct hypothalamic visual response classes in diurnal *Rhabdomys*. A) Mean chirp-evoked response profiles for each functional cluster in *Rhabdomys*, with the stimulus profile shown above. Functional clusters were identified by Gaussian mixture model clustering of sparse principal components derived from chirp-response PSTHs. B) Sequential coronal schematics through the anterior hypothalamus, arranged from rostral to caudal, showing the estimated anatomical locations of recorded *Rhabdomys* units assigned to each cluster. Clusters are shown from top to bottom as clusters 1 - 5. C) Polar plots summarising light-step response polarity and amplitude for units in each cluster. D) Cumulative distribution functions showing sustainedness index distributions across clusters. E, F) Cumulative distribution functions showing the proportion of cells within each cluster with significant responses to E) the temporal-frequency chirp and F) the contrast chirp, based on trial-to-trial response correlations tested against shuffled responses. G) Cumulative distribution functions comparing contrast sensitivity across clusters, quantified as C50 only for responsive units. H) Cumulative distribution functions comparing temporal-frequency tuning across clusters, quantified as Δ amplitude between responses at 2 Hz and 8 Hz.

The cluster mean PSTHs are consistent with categorical inter-cluster differences in visual response characteristics. These were borne out by individual unit level analyses. Polar plots of ON *vs* OFF responses (Fig. 4C) confirmed that clusters 1 and 4 are dominated by excitatory responses to lights on, cluster 2 neurons are excited by both ON and OFF, and clusters 3 and 5 by light OFF. Sustainedness index analysis (Fig. 4D) reveals cluster 2 and 1 as being most transient and cluster 5 the most sustained. The PSTH suggests that cluster 2 units hardly respond to temporal and contrast chirps and, indeed, they showed low trial-to-trial correlation for this stimulus component (Fig. 4E-F). All other clusters had strong responses to both contrast and frequency modulations. Clusters 1 and 3-5 showed linear contrast response relationships with no strong difference in contrast sensitivity (Fig. 4G). These clusters differed in temporal frequency response relationships, with cluster 5 having broadband tuning, while all other clusters were low-pass (Fig. 4H). These data thus reveal that visually responsive units in the *Rhabdomys* anterior hypothalamus fall into at least five functionally distinct clusters differentially distributed across the SCN and peri-SCN (Table 1).

**Table 1.** Cluster characteristic in *Rhabdomys* hypothalamus.

| Cluster | Location | Acute response | Tonic response | Temporal frequency tuning | Contrast |
| --- | --- | --- | --- | --- | --- |
| Cluster 1 | SCN | strong ON | ON | low-pass | linear |
| Cluster 2 | SCN | strong ON | - | - | - |
| Cluster 3 | SCN and lateral peri-SCN | strong ON | OFF | broadband | linear |
| Cluster 4 | SCN | weak ON | ON | low-pass | linear |
| Cluster 5 | rostral peri-SCN | ON and OFF | OFF | broadband | linear |

## Discussion

Retinal innervation of the suprachiasmatic region in mammals is typically considered as a conduit for irradiance information to support synchronisation of the SCN circadian clock with the 24 h light:dark cycle (1). Here, we show that in a diurnal mammal this view under-represents both the richness and anatomical extent of visual information in this part of the brain. Although the canonical SCN response to ambient light reported in nocturnal rodents (2–11) is preserved in *Rhabdomys pumilio*, responses to more dynamic stimuli reveal that the SCN region contains at least five functionally distinct information classes. The neurons conveying this information extend beyond the SCN, especially in rostral and lateral directions. Those peri-SCN neurons differ systematically from SCN neurons in both distribution and response profile, indicating different roles in vision.

Until now assessments of light response in diurnal SCN have encompassed small numbers of neurons presented with a limited array of stimuli (light pulses, sometimes at different intensity) (17, 18). Applying modern recording and analysis technologies has enabled us to record much larger numbers of neurons across a much wider array of visual stimuli. Our data do not support the view that there is a fundamental distinction between the light response in diurnal *vs* nocturnal SCN. Many *Rhabdomys* SCN neurons show sustained responses to light steps and the implication that they could encode ambient light in their maintained firing rate is confirmed by progressive changes in firing in response to the staircase of sequentially increasing and decreasing irradiance (Fig. 2). Although we did find units inhibited by light in *Rhabdomys* SCN, these were in fact no more common (5/83) than we find in *Mus* (11/101). Given the relatively large number of units sampled we can be sure that, at least under these experimental conditions, *Rhabdomys* do not support the hypothesis that the diurnal SCN is reliably more inhibited by light (17, 18). Rather, our data are consistent with a parsimonious interpretation that in both diurnal *Rhabdomys* and nocturnal *Mus* SCN ambient light is encoded according to a monotonic (primarily excitatory) irradiance response relationship (11). This would align well with its essential visual task of measuring ambient light as a signal of time of day in both diurnal and nocturnal species. One might expect diurnal species to show lower sensitivity to this stimulus given their greater exposure to bright daylight, but we do not, and in fact the maintained activity of SCN neurons in both *Mus* and *Rhabdomys* seem to be tuned to track irradiances typical of the twilight transition.

In addition to irradiance-dependent changes in maintained firing, responses to more dynamic visual features (e.g. step onset/offset) were common in *Rhabdomys*. The representation of dynamic visual stimuli is substantially enhanced in *Rhabdomys* compared to *Mus*. Within the SCN itself, acute light response components are larger and have higher sensitivity in *Rhabdomys*, while in the neighbouring hypothalamus visual responses were rare in *Mus* but abundant (63% of recorded units) in *Rhabdomys*. As these peri-SCN responses occur in units lacking the circadian rhythm of SCN neurons and were more biased towards transient response components we can be confident that they represent distinct neuronal populations. The differences between Rhabdomys and Mus thus constitute an expansion of the hypothalamic visual centre associated with diurnality to match that previously described for visual thalamus and colliculus (19, 20, 28). The extent to which this is replicated in other diurnal species remains to be determined, but fMRI and neuroanatomical data are consistent with presence of a diffuse suprachiasmatic visual system in primates similar to that described here (21, 36).

The more dynamic visual response components described here have no obvious utility to encode the daily change in ambient light and, at least in mice, circadian photosensitivity is not impacted by visual contrast (12). The appearance of such robust responses to dynamic stimuli in *Rhabdomys* thus implicates this suprachiasmatic region in visual behaviours beyond establishing daily rhythms (37). A first step to defining hypotheses regarding the function of such dynamic visual responses is a description of the visual features to which these neurons are sensitive. Units in the *Rhabdomys* suprachiasmatic region, show substantial diversity in their response to even the stylised and limited chirp stimulus employed here. We used unsupervised clustering to group units according to their responses across multiple stimulus components and provide an assessment of the number of parallel visual information streams required to capture this diversity. Although the number of clusters identified by this approach depends on the stimuli used, the five clusters found in the *Rhabdomys* suprachiasmatic region showed distinct response properties. Together, they provide a useful framework for understanding the diversity of visual information streams in this part of the brain.

The range of visual response types in *Rhabdomys* suggests a diversity of visual functions. Clusters 1 and 3 - 5 have sustained responses to a light step indicating that they have the capacity to encode ambient light. However, their response to chirp stimuli suggests that while their maintained firing rate may encode scene irradiance finer timing of spikes conveys information about more dynamic visual features. In terms of feature selectivity, cluster 5 showed broader temporal-frequency tuning than other groups, a response property that in other visual circuits is often associated with sensitivity to rapidly changing visual input (38). Whether these neurons contribute to object-motion or behaviourally guided visual responses remains to be tested. Cluster 5 neurons are largely restricted to a region rostral to the SCN, raising the possibility that this is a new visually responsive location contributing to some visually engaged behaviours or their modulation. Clusters 1, 3 and 4 have similar sensitivity to temporal frequency and contrast selectivity but differ in the relative amplitude of responses to these dynamic stimuli *vs* steady light and in response polarity. The low-pass temporal frequency tuning and linear contrast sensitivity of these units could allow them to contribute to pattern detection tasks. Cluster 2 neurons are found in the SCN and diffusely across neighbouring regions and represent outliers insofar as they have limited sustained response to light. This implies that they lack the capacity to encode ambient light. In fact, units in this group respond primarily to light step onset and little to the more dynamic contrast or temporal frequency chirps. These features suggest that these neurons are tuned to detect sudden increases in local or scene brightness. This cluster resembles response motifs described in retinal pathways involved in detecting abrupt luminance changes or object-background discontinuities (39–41).

In summary, these findings place SCN irradiance coding within a broader hypothalamic visual system in a diurnal mammal. Rather than acting solely as a circadian light meter, the suprachiasmatic region of *Rhabdomys* contains distributed neuronal populations capable of encoding both ambient illumination and dynamic visual change. This expanded visual representation provides a potential substrate through which light may shape physiology and behaviour in day-active mammals beyond circadian entrainment alone.

## Material and methods

### Animals and ethical approval

All experiments were conducted in accordance with the UK Animals (Scientific Procedures) Act 1986 and were approved by the Home Office under project licence PP3176367. In total, thirty-one adult *Rhabdomys pumilio* and nine adult *Mus domesticus* (C57BL/6) from the University of Manchester were used. Of the *Rhabdomys*, two animals were used for cholera toxin intravitreal injections (one male and one female, 8 months old), eighteen for the c-Fos experiment (six females and twelve males, 6 - 10 months old), and eleven for electrophysiology (one female and ten males, 4.5 - 11 months old, 61 - 86 g). Nine male *Mus* were used for *in vivo* electrophysiology (4 - 7 months old, 23 - 39 g). Before experimental procedures, animals were group-housed under a strict 12:12 h light/dark cycle at 22°C, with food and water available *ad libitum*.

### Retinal tracing and tissue clearing

Two *Rhabdomys pumilio* received intraocular injections of 5 µl cholera toxin subunit B conjugated to Alexa Fluor 555 (CTb; Thermo Fisher Scientific). Animals were anaesthetised with isoflurane (5% at 0.5 litres/min), and pupils were dilated with topical 1% tropicamide. A drop of Viscotears and a coverslip were placed on the eye to allow direct visualisation of the needle tip under an operating microscope during injection. An ultrafine 30-gauge Hamilton RN needle attached to a 10 µl Hamilton glass syringe was inserted through the scleral equator (*ora serrata*) into the vitreous cavity, taking care to avoid the lens. CTb was injected slowly, after which the needle was withdrawn gradually and topical analgesic was applied (0.25% bupivacaine hydrochloride; Aspen). Animals were perfused 5 days later.

### Perfusion

Animals were anaesthetised with isoflurane (5% at 0.5 litres/minute) in their home cages and maintained with 3 g/kg intraperitoneal injection of urethane (30%, w/v) before they were transcardially perfused using 4% paraformaldehyde (PFA). After perfusion, the brain was carefully harvested and post-fixed overnight in 4% PFA at 4°C.

### CTb visualisation

Immediately following perfusion, eyes and brains were removed and post-fixed in 4% PFA. The retina from the injected eye was carefully dissected from the eye cup, mounted flat and imaged as a wholemount using the Andor Dragonfly inverted confocal microscope equipped with X20 Nikon 20x/0.45 NA S Plan Fluor C. S. Corr objective, to visualise retinal transduction of CTb-555 (Supplementary Fig. 5). After fixation, rostral and caudal regions of the brain were removed leaving a ∼3 mm thick coronal slice incorporating the hypothalamus. This slice was cleared using the iDISCO clearing method (42), adapted without immunolabelling steps. Endogenous fluorescent signal in cleared slices was imaged on the Zeiss Z1 Light Sheet microscope equipped with The ZEISS Clr Plan-Neofluar 20x/1.0 objective. Samples were imaged in ethyl cinnamate (Sigma Aldrich). All image post-processing was done in Imaris 10.1 (Oxford Instruments) image analysis software. Z-stack images were converted into videos and exported using Zeiss Zen (blue edition).

### c-Fos light stimulation

A total of 18 adult *Rhabdomys pumilio* were maintained under a stable 12:12 h light/dark cycle before stimulation. At ZT16, animals were transferred to a test chamber and exposed for 30 min to either steady light or 4 Hz flickering light at 400 melanopic lux EDI or kept in darkness as control. Control animals were kept in darkness for 30 min in the same experimental setup. After stimulation, all animals remained in darkness for an additional 30 min before perfusion, which was performed in the dark.

### c-Fos immunohistochemistry and quantification

After perfusion, brains were removed, post-fixed overnight in 4% PFA at 4°C and cryoprotected in 20% sucrose. After approximately two weeks in sucrose, brains were sectioned coronally at 40 µm on a freezing sledge microtome (Series 8000; Bright Instruments Ltd., Huntingdon, UK). Sections were stored in 30% sucrose at 20°C until immunohistochemical processing.

Free-floating sections spanning the rostral-caudal axis of the anterior hypothalamus, including the SCN, were selected for c-Fos immunohistochemistry. Sections were washed in 0.1 M PBS, incubated for 20 min at room temperature in 1.5% hydrogen peroxide prepared in 0.05% Triton X-100/PBS to quench endogenous peroxidase activity, and washed again in PBS. Sections were then blocked for 60 min at room temperature in 10% normal donkey serum in 0.05% Triton X-100/PBS before incubation for 48 h at 4°C with rabbit anti-c-Fos antibody (1:400; 9F6; Cell Signaling Technology, MA, USA) diluted in 5% normal donkey serum in 0.05% Triton X-100/PBS. After washing, sections were incubated for 90 min at room temperature with biotinylated donkey anti-rabbit IgG (1:400; Jackson ImmunoResearch, PA, USA), followed by avidin–biotin complex (1:200; Vector Laboratories, Peterborough, UK) for 90 min. c-Fos immunoreactivity was visualised using nickel-intensified diaminobenzidine for 20–22 min, producing dark blue-black nuclear staining. Sections were mounted on gelatine-coated slides, dehydrated through graded ethanol, cleared in Histoclear, and coverslipped with DPX mounting medium.

Mounted sections were scanned in bright field at a single focal plane using an automated slide scanner with a 20× objective. Sections were anatomically categorised as rostral, mid, or caudal anterior hypothalamus. c-Fos-positive nuclei were quantified in regions of interest encompassing the SCN and peri-SCN areas. Counting boxes of 0.6 mm^2^ were positioned with reference to the optic tract and third ventricle, such that each box rested on the optic tract and was centred on the third ventricle, extending 500 µm laterally from the third ventricle and 600 µm dorsally from the optic tract (Fig. 1B). Counts were performed blind to lighting condition using QuPath 0.4.3 and ImageJ 1.53q.

After initial counting, heat maps were generated to visualise c-Fos activity across very rostral, rostral, mid, and caudal anterior hypothalamic sections by dividing the counting window into 50 µm × 50 µm squares (Fig. 1C). For statistical comparisons across regions, the SCN was identified using in-house DAPI and AVP immunohistochemistry, and SCN-specific counting boxes were drawn on mid and caudal maps, extending 200 µm laterally from the midline and 250 µm dorsally from the optic tract (Fig. 1B). The density of c-Fos-expressing cells was averaged across all SCN bins from these mid and caudal sections to produce a single SCN value per animal (3, 43). Data were assessed for normality using Shapiro–Wilk and Kolmogorov–Smirnov tests. Depending on data distribution, comparisons were performed using one-way ANOVA followed by Tukey’s multiple-comparison test or Kruskal–Wallis test followed by Dunn’s multiple-comparison test. p values < 0.05 were considered statistically significant.

### In vivo electrophysiology

Animals were removed from their home cages 2 - 3 h before lights off, so that all surgical preparations were performed during the light phase. Animals were anaesthetised by intraperitoneal injection of urethane (1.4 - 1.5 g/kg; 20% or 30% solution in sterile saline for *Mus* and *Rhabdomys*, respectively; Sigma-Aldrich, St Louis, MO, USA). Depth of anaesthesia was verified by the absence of withdrawal and ocular reflexes, and an additional dose of urethane equal to 10 - 20% of the initial dose was administered if required, no earlier than 40 min after the initial dose in *Mus* and 60 min after the initial dose in *Rhabdomys*. Atropine (0.3 mg/kg; Sigma-Aldrich) and saline (0.4 mL for *Mus* and 0.7 mL for *Rhabdomys*) were administered subcutaneously after animals had reached a stable plane of anaesthesia, to reduce mucous secretion and maintain hydration. Throughout the experiment, body temperature was maintained at 37.0 ± 0.5°C using a homeothermic blanket system. To minimise variation in retinal illumination, pupils were dilated with topical 1% tropicamide before visual stimulation. A transparent ophthalmic gel was applied to keep the cornea moist throughout the recording. The animal’s head was secured in a stereotaxic frame using ear and incisor bars. The skull was exposed, and two reference points were identified: *bregma* and *lambda*. Care was taken to level the skull by adjusting the incisor bar so that the bone surfaces near bregma and lambda were at the same relative height, differing by no more than 100 µm. The method used to define bregma and lambda was based on (44), which was particularly important for *Rhabdomys pumilio* recordings because the sutures were often irregular and the intersection between the coronal and sagittal sutures was positioned too far caudally to represent the true *bregma*. *Bregma* was therefore estimated as the intersection between an imaginary parabola fitted to the edges of the coronal suture and a straight midline perpendicular to it, usually aligned with the coronal suture (Supplementary Fig. 6).

Coordinates for targeting the *Mus* SCN were based on the stereotaxic mouse brain atlas of Paxinos and Franklin (31). Craniotomies were made 1.0 mm lateral and 0.1 mm caudal to bregma. All recordings were performed with the probe positioned sagittally at a 9° angle relative to the dorsal–ventral axis. The rostral shank of the recording probe was positioned 0.8 mm lateral and 0.0 mm caudal to bregma, and the electrode was lowered to the level of the SCN, 4.5 - 5.5 mm from the brain surface.

Coordinates for *Rhabdomys pumilio* recordings were established from cadaver experiments (n = 3). Two approaches were tested to target the SCN: vertical probe insertion and insertion at a 9° angle relative to the dorsal-ventral axis, both with sagittal probe orientation. The angled approach was more successful. Craniotomies were therefore made either 0.5 mm lateral and 0.0 mm caudal to bregma for vertical recordings, or 1.2 mm lateral and 0.0 mm caudal to bregma for angled recordings. Coordinates for the rostral shank were 0.2 - 0.5 mm lateral, 0.4 mm caudal to bregma, and 5.5-6.5 mm below the brain surface for vertical insertions, and 1.0 - 1.3 mm lateral, 0.4 mm caudal to bregma, and 6.0 - 7.3 mm below the brain surface for angled insertions.

A Buzsáki 32L probe (Neuronexus, Ann Arbor, MI, USA), consisting of four shanks spaced 200 µm apart, was used for extracellular recordings. Before the first insertion, the recording probe was dipped in the fluorescent dye CM-DiI (V22888; Thermo Fisher Scientific, Waltham, MA, USA) to enable subsequent histological verification of recording placements. A fluid-filled micromanipulator (MO-10; Narishige International Ltd, London, UK) was used to move the probe along the dorsoventral axis.

Recording placements were identified electrophysiologically based on cellular responses to tested light stimulation. Once light-induced activity was detected, animals were dark-adapted for 30 min to allow neuronal activity to stabilise. Neuronal activity was acquired using a Recorder64 system (Plexon Inc., Dallas, TX, USA), amplified ×3000, high-pass filtered at 300 Hz, and digitised at 40 kHz. After each recording, the probe was either advanced deeper or slowly retracted and moved laterally or medially to sample additional units from non-overlapping regions.

### Histological reconstruction of recording sites

After completion of all recording protocols, animals were intracardially perfused with saline followed by 4% PFA. Brains were removed, postfixed overnight at 4°C and cryoprotected using 30% sucrose in phosphate-buffered saline (PBS). Freezing microtome was used to prepare coronal slices (40 µm). Immunofluorescence labelling for arginine vasopressin (AVP) and/or DAPI was performed to visualise anatomical boundaries of the SCN. Briefly, following washes in 0.1 m PBS and 0.1% Triton X-100 in PBS, sections were blocked for 1 h in 5% normal donkey serum. Then, they were incubated with primary antibody (dilution 1:5000; AVP Rabbit; AB1565; Millipore, Burlington, MA, USA) for 2 days at 4°C. After washing, sections were incubated with Alexa Fluor 488 Donkey anti-Rabbit IgG (dilution 1:800; A-21 206; Invitrogen, Thermo Fisher Scientific) overnight at 4°C. Finally, sections were mounted onto gelatine coated slides and cover-slipped using ProLong Gold Antifade Mountant (Molecular Probes, Invitrogen; Thermo Fisher Scientific). Photos were taken using a DFC365 FX camera (Leica, Wetzlar, Germany) connected to a DM2500 microscope (Leica) with LAS AF6000 software (Leica).

Recording locations were reconstructed by aligning stereotaxic records, probe depth, shank geometry, CM-DiI-labelled electrode tracks, and AVP and DAPI immunolabelling of the SCN. For each recording placement, the estimated position of each shank and recording site was projected onto the nearest matched coronal section using the Paxinos and Franklin mouse brain atlas as a reference and species-specific *Rhabdomys* SCN anatomy from AVP immunohistochemistry (34), as well as densely packed DAPI staining. The SCN was defined anatomically as the AVP-positive nucleus immediately dorsal to the optic chiasm and adjacent to the third ventricle. Units were classified as SCN if their estimated recording position overlapped this AVP-defined boundary. Units were classified as peri-SCN if their estimated position was outside this boundary, either rostral to the SCN or lateral to its border, within the anterior hypothalamic region sampled by the probe. These anatomical assignments were made before light-response properties were analysed.

### Visual stimulation

A pE-4000 system (CoolLED, Andover, UK) connected to a liquid light guide fitted with a diffuser (Edmund Optics, Barrington, NJ, USA) was used as a light source. The diffuser was positioned at ∼5 mm from eye contralateral (or ipsilateral for some recordings) to the recording site. White light was used for all stimuli (Supplementary Figure 7), consisting of output from four LEDs with peak emission at 385, 460, 550 and 635 nm. Irradiance was measured by SpectroCAL MKII spectroradiometer (Cambridge Research System, Cambridge, UK) and all irradiances are presented as species specific melanopic equivalent daylight illuminance (melanopic lux EDI) (45). Three main light protocols were used (in different combinations, see Supplementary Table 2 for numbers): full-field light step of increasing irradiance, staircase stimuli and full-field chirp. Initial light test used to classify cells as light-responsive was 10 s long bright full-field light step (5789 melanopic lux EDI) presented 10 times with an interval of 50 s. Neutral density filters (Thorlabs Inc., Newton, NJ, USA) were used to produce increasing range 0.005 - 5789 melanopic lux EDI. Staircase stimulus was described in detail previously (13). In short, it consisted of six ascending and following descending steps in the range 0.002 – 2449 melanopic lux EDI controlled by motorised filter wheel (FW102C/FW212C Series; Thorlabs Inc., Newton, NJ, USA). Such pyramid was repeated 3 -8 times. Each step consisted of 30 s of steady light, followed by 16 s of superimposed temporal WN, 8 s of temporal chirp, and finished with 8 s of contrast chirp. Full-field chirp was presented at 258.88 melanopic lux EDI and was repeated 9-10 times. The chirp comprised of a 3 s light step from dark followed by sinusoidal modulations of increasing temporal frequency from 1 to 8 Hz at 1 Hz/s speed at 98% Michelson contrast from photopic background and sinusoidal contrast modulation at 2 Hz increasing from 3% to 97%. 2 s of steady light preceded temporal and contrast elements of the chirp.

### Spike sorting

Spike sorting was performed offline on high-pass-filtered extracellular recordings. Automated spike detection and initial clustering were conducted using Kilosort2 (46). Identified clusters were then exported to OfflineSorter (Plexon Inc., Dallas, TX, USA) as virtual tetrodes, using spike waveforms detected across four adjacent channels, and manually curated. Only well-isolated single units were included in subsequent analyses. Unit isolation was assessed using waveform shape, cluster separation, multivariate analysis of variance F statistics, J3 and Davies-Bouldin validity metrics, and interspike interval distributions. Units were accepted only when interspike interval histograms showed a clear refractory period with minimal violations within 1 ms. Multiunit activity and unstable clusters were excluded. To minimise duplicate sampling, recording placements were treated as independent only when probe position was changed to a non-overlapping region based on micromanipulator displacement, probe geometry and post hoc histological reconstruction.

### Statistical analysis

In multiunit recordings from 22 placements in *Rhabdomys* (n = 11), we identified 362 single units from which 147 were located within the borders of the SCN. In parallel *Mus* recordings 383 units were identified from 20 placements (n = 9) and 167 were estimated to fall within the boundaries of the SCN.

### Identifying light-responsive units

To identify light-responsive cells, 10 repeats of 10 s long full-field bright light step were presented with 50 s inter-stimulus interval. We classified cells as light-responsive (LR) if the change in their firing rate during light stimulation, either peak onset (0 - 300 ms), tonic part (5 - 10 s after light onset) or peak offset (0 - 300 ms after light offset) was ± 2 SD of their baseline firing recorded in the dark just before light stimulations.

### Response polarity analysis

Response polarity was quantified from light-step responses by comparing changes in firing around light onset and light offset. For each unit, spikes were counted in 1 s windows immediately before and after stimulus onset and offset for each trial. The mean spike count across trials was then calculated for each window, and changes in firing were expressed as the difference between the post-stimulus and pre-stimulus windows for light onset (ΔON) and light offset (ΔOFF).

Each unit was represented in Cartesian coordinates using ΔON and ΔOFF values and these coordinates were converted to polar form. The resulting vector angle was used as a measure of response polarity, whereas vector magnitude represented overall response amplitude. Kuiper test was used to compare between distributions of vector angle.

### Sustainedness index

The sustainedness index (based on (47)) was used to quantify the extent to which light-evoked firing was maintained throughout the stimulus rather than confined to transient onset responses. For 10 s light-step responses, PSTHs (peristimulus time histograms) were generated across a 15 s window comprising 5 s before and 10 s during light presentation.

Responses were min-max normalised within the analysis window. The mean firing rate before and after light onset was compared to determine response polarity. For units showing an increase in firing after light onset, the area under the normalised response curve during the light step was calculated and divided by the number of bins. For units showing a decrease in firing after light onset, the normalised response was inverted before calculating the area under the curve. This yielded an index ranging from 0, indicating highly transient responses, to 1, indicating highly sustained responses. Distributions were compared using two-sample Kolmogorov–Smirnov tests, with results reported as p values and D statistics.

### Irradiance coding

For full-field light steps data, PSTHs were generated (only LR cells excited by light were used) using 100 ms bins from 5 s before light onset to 15 s after light onset, encompassing the full 10 s light step. Changes in firing rate were calculated for the acute onset response, defined as the first 300 ms after stimulus onset, and for the tonic response, defined as 5 - 10 s after stimulus onset. Irradiance-response relationships were fitted with a four-parameter variable-slope function, with the lower asymptote constrained to zero. Differences in fitted irradiance-response relationships between conditions were assessed using F tests comparing EC50, Hill slope and/or maximal response.

Staircase stimulus was analysed as previously described (13). In short, LR units firing rate was averaged separately during steady-light, white-noise and chirp epochs at each irradiance step, then normalised to each unit’s maximum response across the full staircase protocol. Only stable single units maintained across at least four staircase presentations were included. Baseline-subtracted, normalised firing rates were fitted with four-parameter sigmoidal irradiance-response curves, with the lower asymptote constrained to zero. Differences in fitted irradiance-response relationships between conditions were assessed using F tests comparing EC50, Hill slope and/or maximal response.

### Identifying chirp-responsive units

To identify units responsive to the chirp stimulus, PSTHs were generated using 25 ms bins from 2 s before onset of the initial 3 s chirp step to 32 s after step offset, giving a total analysis window of 34 s. Responsive units were identified by comparing trial-to-trial correlation across the full chirp response with a null distribution generated by shuffled-bin analysis. The null distribution was generated from 500 repeats of shuffled data, and units were classified as chirp responsive when p < 0.001. Units were required to contain spikes in at least eight trials to be considered responsive.

Trial-to-trial correlation was defined as the correlation of response profiles across repeated stimulus presentations.

### Unsupervised clustering of chirp responses

Functional clustering was performed on the pooled dataset of *Rhabdomys* and *Mus* units from all sampled regions that showed significant responses to the full chirp stimulus. Mean chirp response PSTHs were generated using 50 ms bins and used to compute sparse principal components.

Thirty-seven sparse principal components were extracted and used as inputs for Gaussian mixture model clustering. The Gaussian mixture model was run with random initialisation, and the optimal number of clusters was selected using Bayesian model comparison, based on the lowest Bayesian information criterion and a Bayes factor threshold of < 6.

The distribution of units across clusters was compared between groups using a shuffle test based on the relative proportion of units assigned to each cluster.

### Sustained Index

The sustained index was calculated as for the 10 s light step, except that different time bins were used for 10 s step and chirp step analyses.

For the chirp step, responses were analysed in a 5 s window comprising 2 s before and 3 s during the step. Responses were min-max normalised within each analysis window. The mean firing rate before and after step onset was then compared. If firing increased after step onset, the area under the normalised response curve was calculated during the step and divided by the number of bins. If firing decreased after step onset, the normalised response was inverted and the area under the inverted curve was calculated during the step and divided by the number of bins.

### Contrast chirp analysis

Contrast responses were analysed from chirp stimulus trials using peristimulus time histograms generated with 25 ms bins from 2 s before chirp step onset to 30 s after chirp step offset.

Contrast response amplitude was calculated for each contrast modulation as the peak-to-trough firing-rate difference. Baseline amplitude was calculated over an equivalent 0.5 s window during steady light immediately before contrast chirp onset and subtracted from each contrast response amplitude.

For units with monotonic contrast responses, contrast sensitivity was estimated by fitting a four-parameter Naka-Rushton function using least-squares minimisation in MATLAB (*lsqcurvefit*). The fitted parameters were minimum response, maximum response, C50 and slope. Parameter constraints were set using the observed response range: minimum response was constrained between the minimum and maximum measured amplitude, maximum response was constrained between the maximum measured amplitude and 10% above that value, slope was constrained between 0.01 and 5, and C50 was constrained between 0 and 1.

Goodness of fit was assessed by the coefficient of determination (R²). Units were classified as having simple unimodal contrast responses if they were responsive to the contrast chirp, had R² > 0.4, and showed a positive mean normalised amplitude at high contrasts (75-100%). Units responsive to the contrast chirp but failing these criteria were classified as having complex contrast responses. C50 values were extracted only from simple contrast-responsive units.

### Temporal chirp analysis

Temporal-frequency responses were analysed from the same chirp stimulus trials using 25 ms binned peristimulus time histograms.

Temporal response amplitude was calculated for each temporal modulation as the peak-to-trough firing-rate difference. Amplitudes were averaged across equivalent temporal-frequency repeats to obtain eight frequency-response values per unit. Baseline amplitude was calculated during a steady light window immediately before chirp onset and subtracted from each response amplitude.

Temporal-frequency tuning was summarised by calculating the difference between normalised response amplitude at 2 Hz and 8 Hz (Δ amplitude, 2 Hz minus 8 Hz). Positive values indicated stronger responses at lower temporal frequencies, whereas values close to zero indicated broader temporal-frequency tuning.

## Acknowledgements

This study was funded by European Union’s Horizon 2020 research and innovation programme under the Marie Skłodowska-Curie grant (897951) to PO-F, Wellcome Trust Investigator Award (210684/Z/18/Z) to RJL and Wellcome Trust Discovery Award (321693/Z/24/Z) to RJL, TMB and BB-O. AEA was supported by a Sir Henry Dale Fellowship (218556/Z/19/Z), jointly funded by the Wellcome Trust and the Royal Society, TMB was supported by Leverhulme trust grant (RPG-2022-168), RS was supported by Sir Henry Dale fellowship from Wellcome Trust (220163/Z/20/Z) and a research grant by Biotechnology and Biological Sciences Research Council (BB/V009680/1) and NM was supported by Academy of Medical Sciences grant (SBF008/1071).

We would also like to acknowledge the Bioimaging Facility at the University of Manchester for providing access to microscopes, which was instrumental in the completion of this research. The microscopes used in this study were purchased with grants from the Biotechnology and Biological Sciences Research Council, Wellcome, and the University of Manchester Strategic Fund.

## Author contribution

PO-F and RJL supervised the project; PO-F, BB-O and RJL designed the experiments; PO-F performed all electrophysiological data collection and analysis, with contributions from JR, AEA and RS; BB-O, MD and PO-F performed immunohistochemistry for the electrophysiological experiments; RR, with contributions from PO-F, performed the intravitreal CTb injections and CTb imaging; BB-O and ADA collected, imaged and analysed the c-Fos data, with contributions from PO-F, MD, NM and TMB; PO-F, RJL and AEA designed the light stimuli; FPM provided tools for the electrophysiological experiments; PO-F and RJL wrote the manuscript with input and approval from all authors.

## Supplementary materials

**Supplementary Fig. 1.**
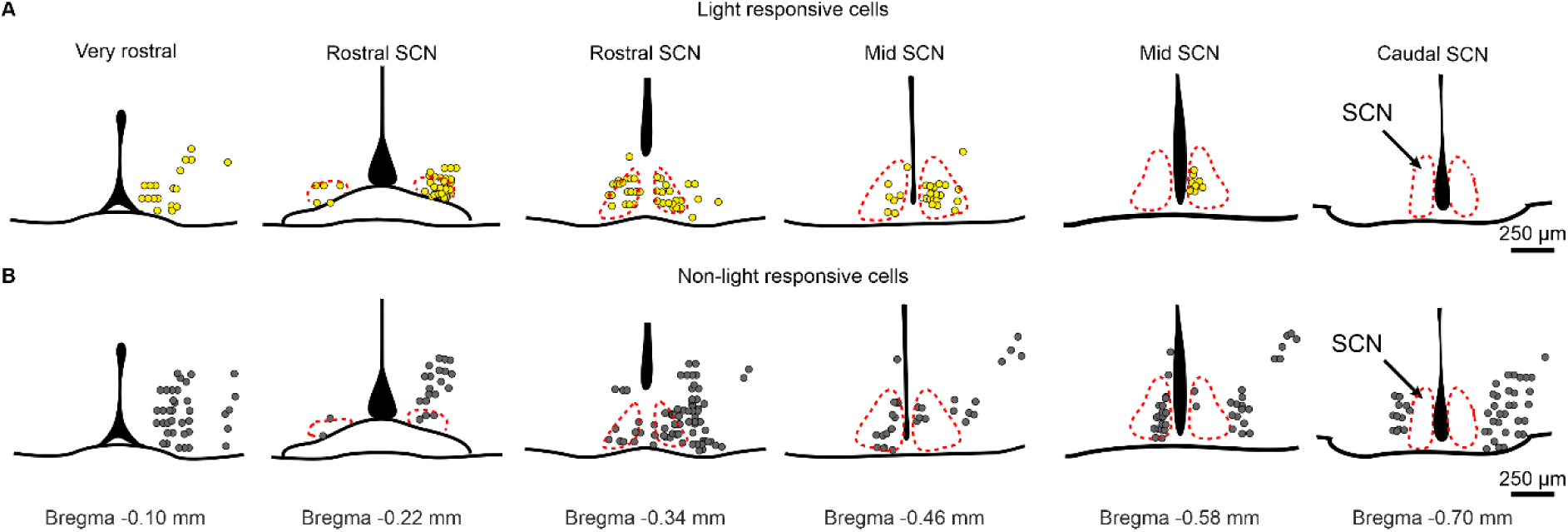
Light-responsive cells are mostly localised to the SCN in nocturnal *Mus.* A, B) Sequential coronal schematics through the anterior hypothalamus of *Mus*, arranged from rostral to caudal, showing the estimated locations of recorded A) light-responsive units and B) non-light-responsive units. Light-responsive units were concentrated within or near the SCN, whereas non-light-responsive units were distributed more broadly across the sampled anterior hypothalamus. Coronal drawings were based on the stereotaxic mouse atlas of Paxinos and Franklin (31).

**Supplementary Fig. 2.**
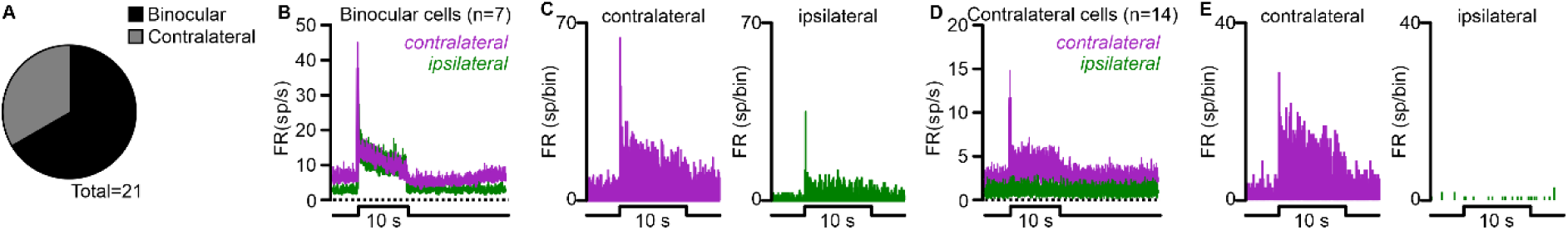
Binocular retinal input into *Rhabdomys* SCN. A) Pie chart summarising the proportion of SCN units responsive to contralateral and/or ipsilateral light stimulation in *Rhabdomys* (n = 21 light-responsive cells from 4 recording placements in 2 animals). Units responding to both contralateral and ipsilateral stimulation were classified as binocular. B) Mean PSTHs ± SEM (bin size = 0.1 s) in response to a 10 s white-light pulse (5789 melanopic lux EDI) for binocularly responsive SCN units, shown separately for contralateral stimulation (purple) and ipsilateral stimulation (green). C) PSTHs (bin size = 0.1 ms) for a representative binocular SCN unit responding to contralateral stimulation (left, purple) and ipsilateral stimulation (right, green). D) Mean PSTHs ± SEM (bin size = 0.1 s) in response to a 10 s white-light pulse for SCN units responsive only to contralateral stimulation, shown separately for contralateral stimulation (purple) and ipsilateral stimulation (green). E) PSTHs (bin size = 0.1 ms) for a representative contralateral-only SCN unit responding to contralateral stimulation (left) but not ipsilateral stimulation (right).

**Supplementary Fig. 3.**
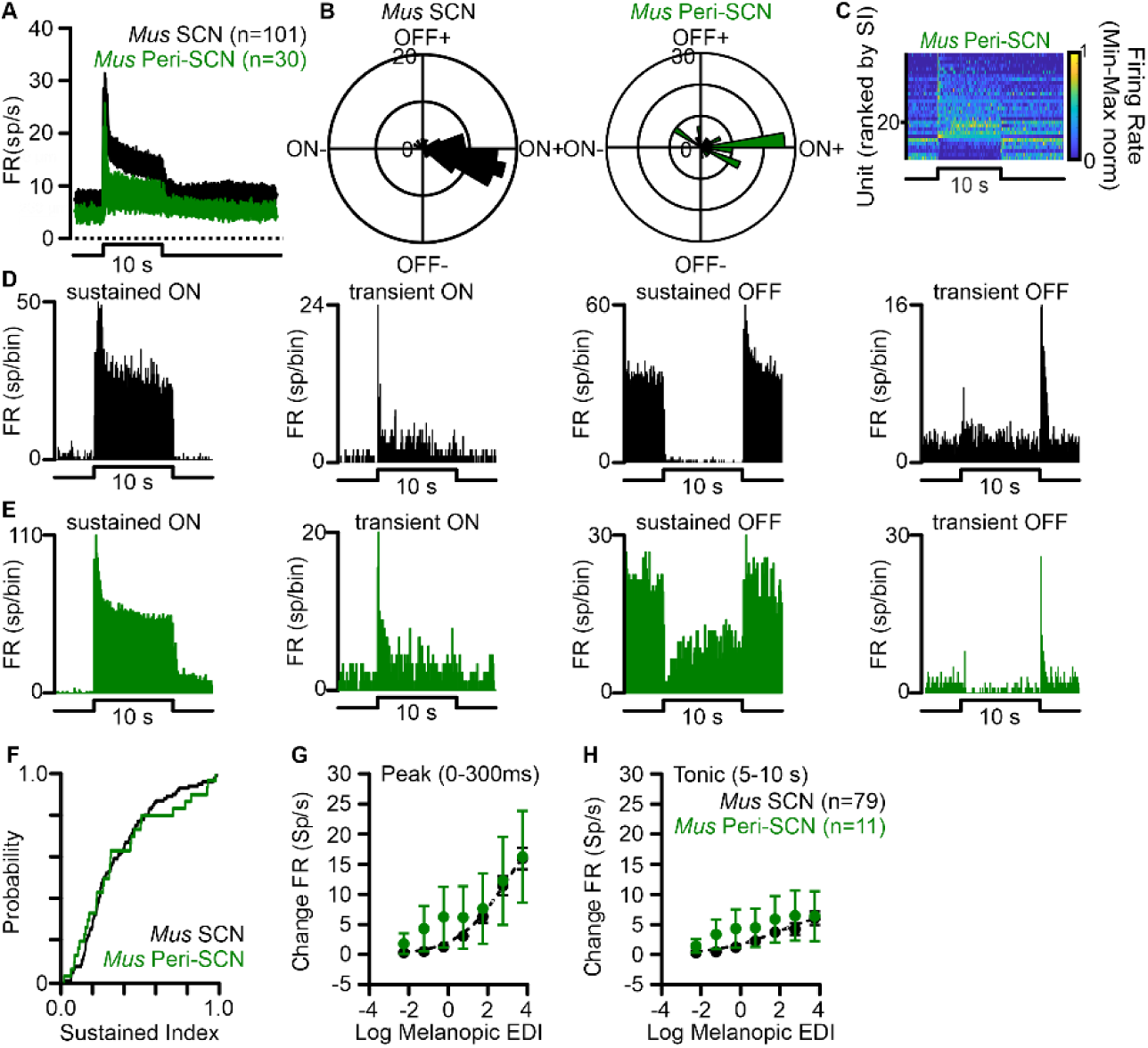
Light-responsive peri-SCN units in nocturnal *Mus* show response properties similar to SCN units. A) Mean PSTHs ± SEM (bin size = 0.1 s) in response to a 10 s white-light pulse (5789 melanopic lux EDI) for all light-responsive SCN units (black, n = 101) and peri-SCN units (green, n = 30) in *Mus*. B) Polar plots summarising response polarity and amplitude show no significant difference between SCN and peri-SCN units (Kuiper test, p = 0.100). C) Heat map of responses to the 10 s light pulse for all light-responsive peri-SCN units in *Mus*, ranked by sustainedness index, with the most sustained responses at the bottom. D, E) PSTHs (bin size = 0.1 ms) for representative light-responsive D) SCN and E) peri-SCN units in *Mus*. F) Cumulative distribution functions comparing sustainedness index distributions between SCN and peri-SCN units show no significant difference between regions (two-sample Kolmogorov–Smirnov test, p = 0. 8667, D = 0.1208). G, H) Irradiance-response relationships for peak firing-rate changes G) during the first 300 ms after light onset (F test, p = 0.2363, F = 1.418) and H) between 5 and 10 s after light onset (F test, p = 0.0659, F = 2.411).

**Supplementary Fig. 4.**
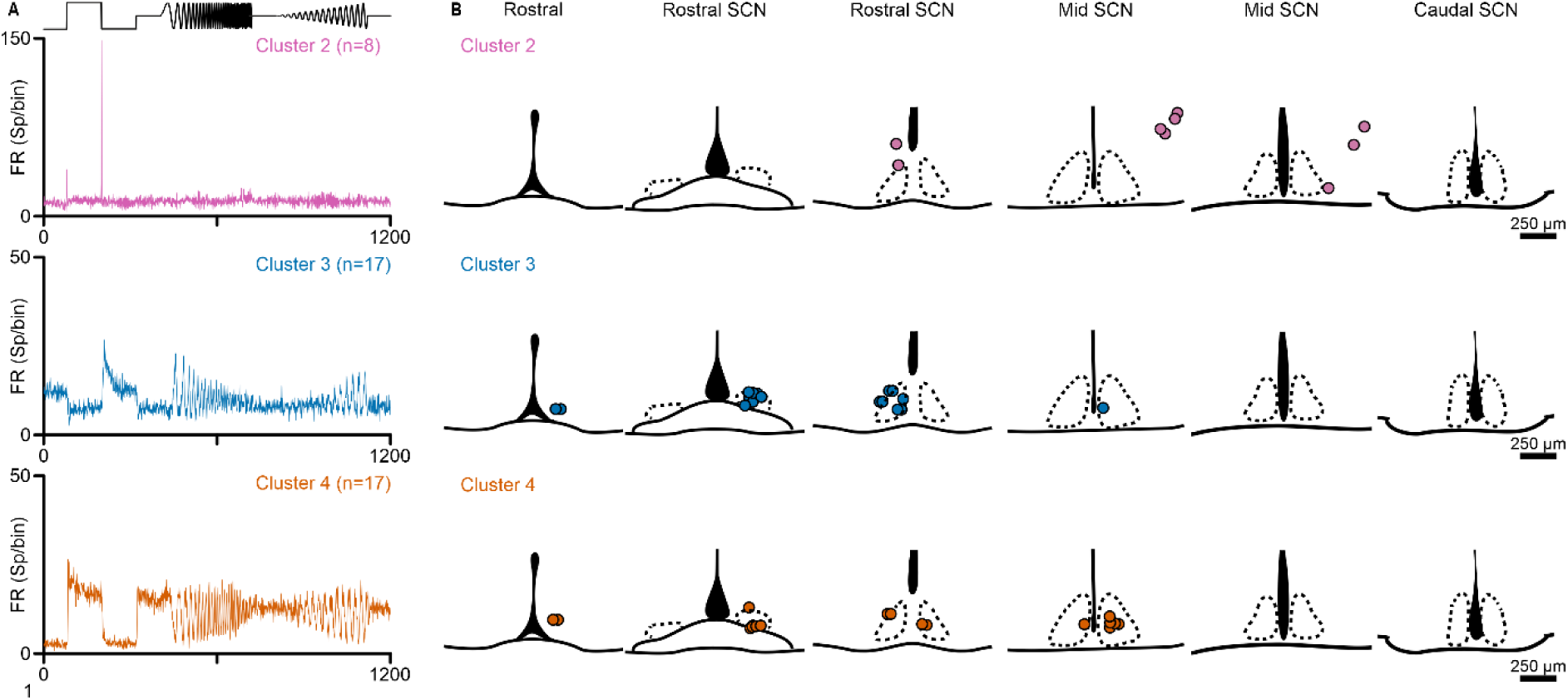
Functional clustering of light-responsive hypothalamic units in nocturnal *Mus*. A) Mean chirp-evoked response profiles for *Mus* units assigned to functional clusters, with the stimulus profile shown above. Clusters correspond to those identified by unsupervised clustering of the combined *Rhabdomys* and *Mus* chirp-responsive dataset. B) Sequential coronal schematics through the anterior hypothalamus of *Mus*, arranged from rostral to caudal, showing the estimated anatomical locations of recorded units assigned to each functional cluster. Only clusters 2–4 were represented in *Mus*; no *Mus* units were assigned to clusters 1 or 5.

**Supplementary Fig. 5.**
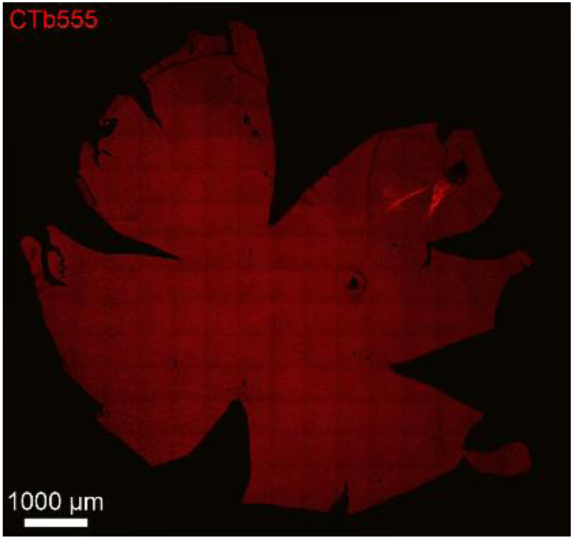
CTb labelling in a flat-mounted *Rhabdomys* retina. Representative flat-mounted *Rhabdomys* retina showing retinal uptake of Alexa Fluor™ 555-conjugated cholera toxin subunit B (CTb-555) following intravitreal injection. The image was acquired using an Andor Dragonfly inverted confocal microscope equipped with a Nikon 20×/0.45 NA S Plan Fluor objective.

**Supplementary Fig. 6.**
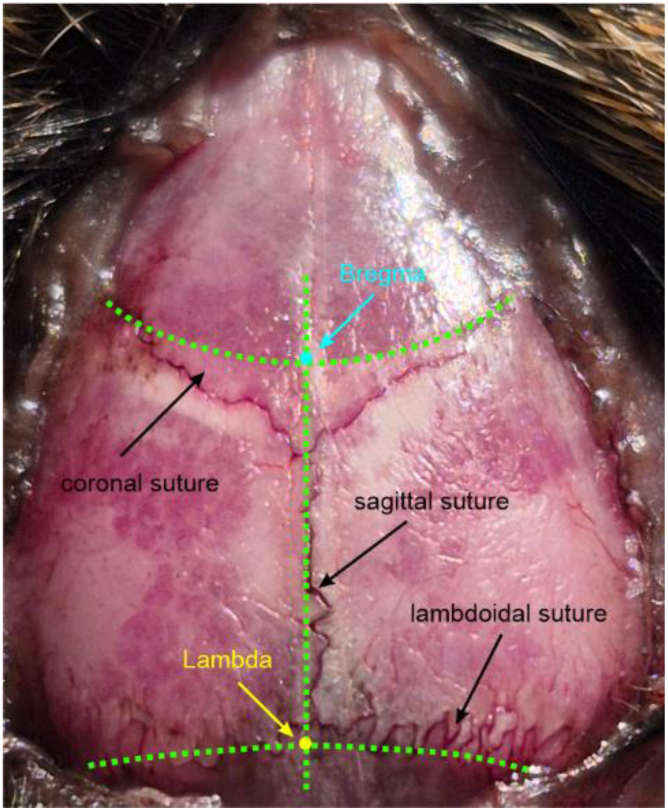
Stereotaxic estimation of *bregma* in *Rhabdomys pumilio*. Representative photograph of the skull surface in *Rhabdomys pumilio*, showing the approach used to estimate the positions of *bregma* and *lambda* for stereotaxic targeting. Drawn guidelines indicate the inferred midline and the estimated intersections used to define these landmarks. The distance between *bregma* and *lambda* was approximately 6.5 mm in the animals used for electrophysiological recordings.

**Supplementary Fig. 7.**
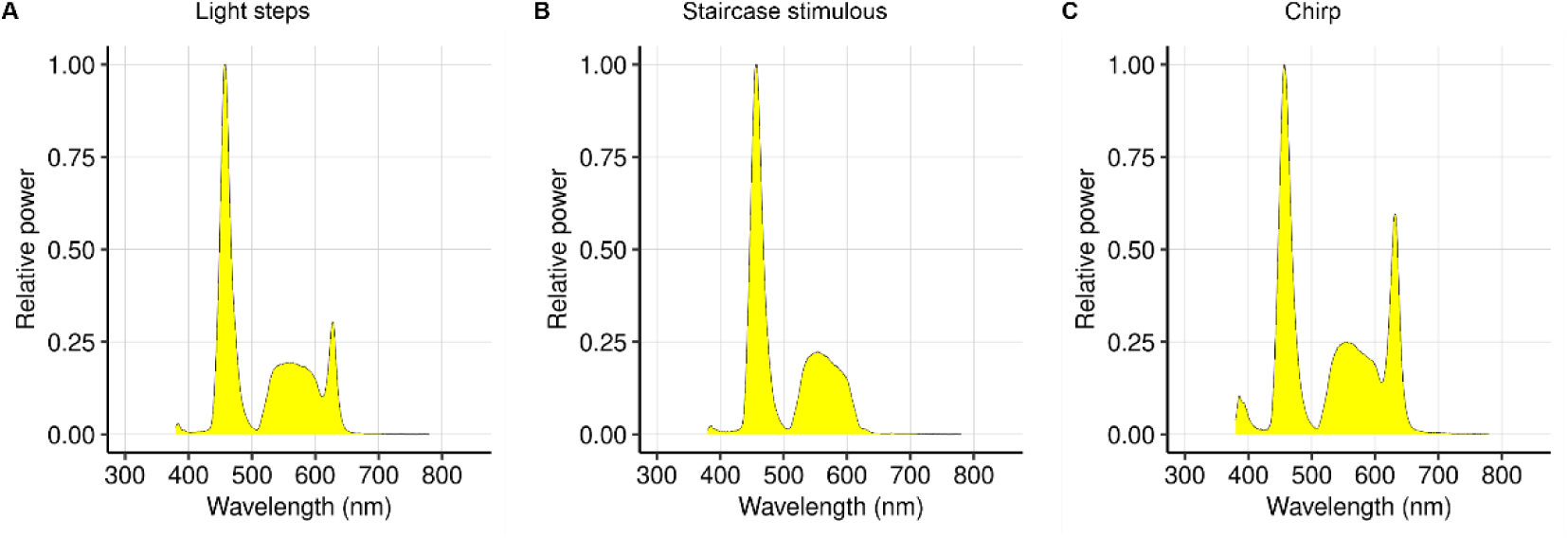
Spectral composition of light stimuli used for visual stimulation. Spectral power distributions of the white-light stimuli generated using a pE-4000 system (CoolLED, Andover, UK). White light was produced by combining output from four LEDs with peak emissions at 385, 460, 550 and 635 nm. Spectra are shown for A) light-step stimuli, B) staircase stimuli and C) chirp stimuli.

**Supplementary Table 1.**
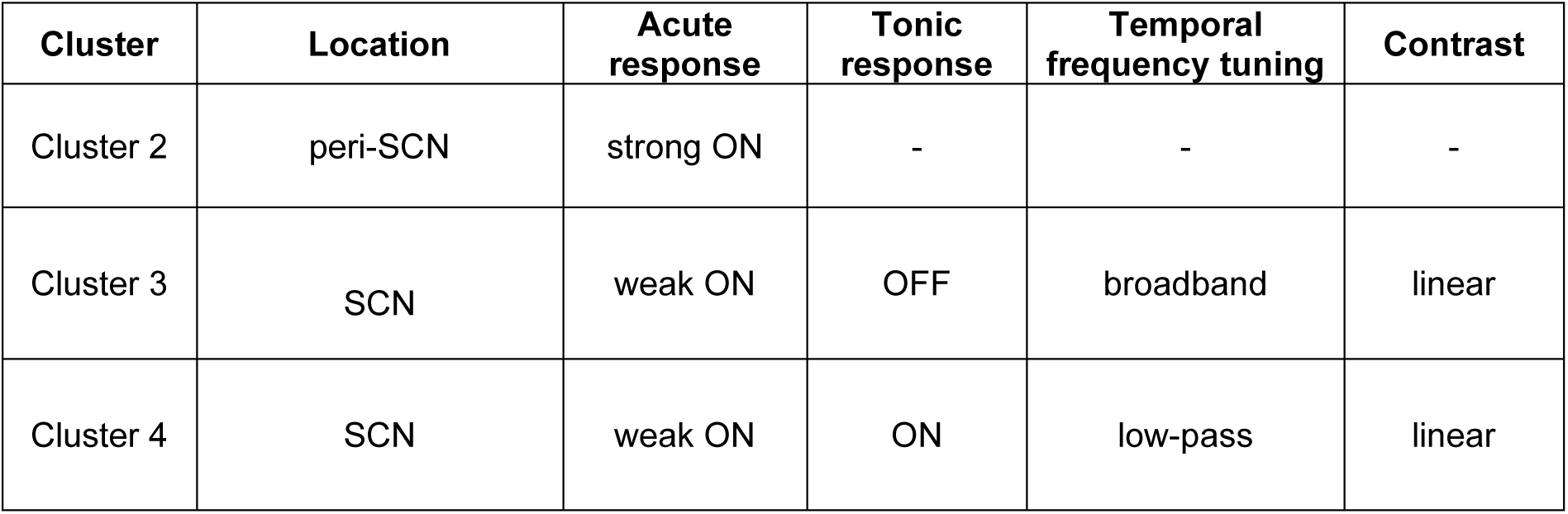
Cluster characteristic in *Mus* hypothalamus.

| Cluster | Location | Acute response | Tonic response | Temporal frequency tuning | Contrast |
| --- | --- | --- | --- | --- | --- |
| Cluster 2 | peri-SCN | strong ON | - | - | - |
| Cluster 3 | SCN | weak ON | OFF | broadband | linear |
| Cluster 4 | SCN | weak ON | ON | low-pass | linear |

**Supplementary Table 2.**
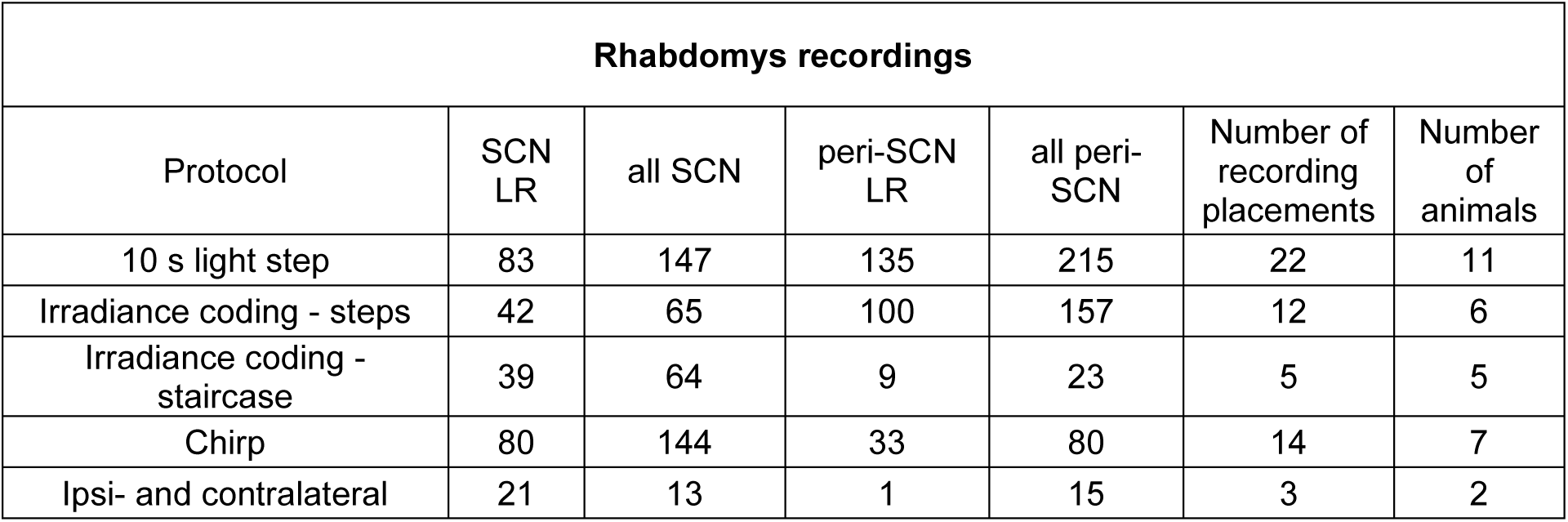
*Rhabdomys* recordings – cells tested by protocol. LR indicates light-step responsiveness, not responsiveness to tested protocol stimuli.

| **Rhabdomys recordings** |  |  |  |  |  |  |
| --- | --- | --- | --- | --- | --- | --- |
| Protocol | SCN<br>LR | all SCN | peri-SCN<br>LR | all peri-SCN | Number of<br>recording<br>placements | Number<br>of<br>animals |
| 10 s light step | 83 | 147 | 135 | 215 | 22 | 11 |
| Irradiance coding - steps | 42 | 65 | 100 | 157 | 12 | 6 |
| Irradiance coding -<br>staircase | 39 | 64 | 9 | 23 | 5 | 5 |
| Chirp | 80 | 144 | 33 | 80 | 14 | 7 |
| Ipsi- and contralateral | 21 | 13 | 1 | 15 | 3 | 2 |

## Notes

### Competing Interest Statement

The authors have declared no competing interest.

